# Do commercial nitrogen-fixing and biostimulant inputs add agronomic value over standard fertilization? An equivalence-based re-analysis of three randomized field trials in the Brazilian Cerrado

**DOI:** 10.64898/2026.06.16.732230

**Authors:** Vitor Horita, Leandro Yuan

**Affiliations:** Independent researcher, North Miami Beach, FL, USA (formerly Chief Agronomist, Horita Group, Western Bahia, Brazil); Cultivo Pesquisa e Desenvolvimento Ltda, São Desidério, BA, Brazil

**Author notes:** Correspondence: Leandro Yuan.

**Keywords:** biological nitrogen fixation, *Azospirillum brasilense*, *Methylobacterium symbioticum*, biostimulants, equivalence testing, statistical power, on-farm trials, Cerrado, plant-microbe inter-actions

## Abstract

Commercial inoculants based on associative diazotrophs and methylotrophic bacteria, and biostimulant programs, are marketed for broad use in cereal and fiber production, yet most field deployment occurs over already adequate mineral fertilization, where their marginal value is poorly quantified. We reanalyzed three randomized complete-block trials conducted at commercial scale in the Brazilian Cerrado (São Desidério, Bahia) under full conventional fertilization: corn (seven treatments, four blocks; 2023/24), cotton (five treatments, four blocks; 2024/25), and soybean (four treatments, six blocks; 2024/25). Treatments evaluated *Azospirillum brasilense, Methylobacterium symbioticum, Bradyrhizobium* spp., a *Bacillus* phosphorus solubilizer, and biostimulants, applied via seed and foliar routes. Beyond conventional analysis of variance, we fit mixed models with block as a random effect, quantified effect sizes and coefficients of variation, computed the minimum detectable difference at 80% power, and applied two one-sided equivalence tests (TOST) against the untreated control at margins of plus or minus 10% and 15% of control yield. No treatment produced a statistically significant yield gain in any crop (all p greater than 0.8; fixed and mixed models concordant). In cotton, the best-powered trial (coefficient of variation 6%), all treatments were statistically equivalent to the untreated control within a 10% margin, an affirmative negative result. In corn and soybean the trials were underpowered (minimum detectable difference 21 to 26% of control), so non-significance is not equivalence; corn showed non-significant numerical gains up to 8.9% with phosphorus-solubilizer and methylotroph combinations that could not be excluded. Foliar nutrient concentrations (single composite per treatment, descriptive) showed no enrichment in inoculated treatments. Under standard fertilization, these commercial inputs delivered no detectable agronomic value where the data were adequately powered to test it. The results clarify the low empirical bar that current associative-fixation products meet in the field and, by extension, the agronomic threshold that engineered, plant-controlled nitrogen fixation must exceed to be useful.

## 1. Introduction

Mineral nitrogen is the largest variable input in cereal and fiber production and a dominant contributor to the environmental footprint of agriculture. The prospect of substituting part of that input with biological nitrogen fixation has driven sustained commercial and scientific interest in microbial inoculants for non-legume crops. Two product classes have reached large-scale field use. Associative diazotrophs such as *Azospirillum brasilense* colonize the rhizosphere and root surface of grasses and can promote growth through a combination of fixation, phytohormone production, and improved root architecture. More recently, the leaf-colonizing methylotroph *Methylobacterium symbioticum*, delivered as a foliar inoculant, has been marketed as a season-long in-planta source of fixed nitrogen across diverse crops. Phosphorus-solubilizing *Bacillus* inoculants and plant biostimulants are frequently stacked with these products in commercial programs.

The agronomic evidence for these inputs is uneven and strongly context dependent. For *Azospirillum brasilense* in maize, a meta-analysis of more than one hundred Brazilian field trials concluded that inoculation can support reductions of roughly 20 to 30% in nitrogen fertilizer without yield penalty, with benefits attributed substantially to root stimulation rather than to fixation alone, and with seed inoculation outperforming foliar application (Barbosa and colleagues, 2022). For *Methylobacterium symbioticum*, controlled evaluations have reported limited and inconsistent effects: a two-year field and pot study in maize found that foliar inoculation did not significantly increase dry matter yield and that estimated fixation was small relative to crop nitrogen removal, leading the authors to question whether the contribution is quantitatively relevant for a commercial product (Rodrigues and colleagues, 2024). A common thread is that demonstrable benefits tend to appear under nitrogen limitation or stress, whereas value added on top of already adequate fertilization is rarely quantified, even though that is the setting in which much commercial deployment occurs.

This question has acquired new significance from the opposite end of the technology spectrum. Engineering nitrogen fixation into cereal-associated bacteria, with nitrogenase expression placed under agriculturally relevant plant or environmental control, has advanced from concept toward field-relevant prototypes (Ryu and colleagues, 2020; Haskett and colleagues, 2022). The promise of such systems is precisely the delivery of agronomically meaningful nitrogen fluxes that present commercial inoculants are not reliably shown to provide. Establishing what current products actually deliver in the field, under realistic management, therefore defines both the unmet need and the bar that engineered solutions must clear.

Most on-farm evaluations report only the outcome of a null hypothesis significance test. A non-significant analysis of variance is widely, and incorrectly, read as evidence that a product has no effect. Non-significance can equally reflect an underpowered experiment that could not have detected an agronomically meaningful difference in the first place. The appropriate tools to distinguish these cases, retrospective power analysis and equivalence testing, are seldom applied to commercial-scale agronomic trials. Here we reanalyze three randomized complete-block trials conducted under full conventional fertilization in the Brazilian Cerrado, asking a deliberately operational question: do commercial nitrogen-fixing and biostimulant inputs add measurable agronomic value over standard fertilization? We pair conventional and mixed-model analysis of variance with effect sizes, minimum detectable difference, and two one-sided equivalence testing, so that, where the data permit, we can state not merely that no difference was detected but whether the treatments are statistically equivalent to the untreated control.

## 2. Materials and Methods

### 2.1 Site and seasons

The three trials were conducted at the agronomic research center of Cultivo Pesquisa e Desenvolvimento at Fazenda Acalanto, São Desidério, Bahia, Brazil (12 degrees 57 minutes 52.6 seconds South, 45 degrees 58 minutes 58.8 seconds West, 833 m elevation), in the rainfed Western Bahia frontier of the Cerrado biome. Corn was grown in the 2023/24 season (emergence 9 January 2024), cotton and soybean in the 2024/25 season (cotton and soybean emergence in late December 2024). On-site meteorological stations recorded temperature, relative humidity, and rainfall throughout each cycle. The corn season was affected by El Niño heat and intermittent water stress; the soybean season included a pronounced 37-day dry spell delivering only 12.2 mm of rain between roughly 49 and 87 days after emergence, coinciding with the reproductive and grain-filling period and depressing yield potential.

### 2.2 Soil and fertilization

Soils were sampled at 0 to 20 cm and characterized as typical clay-loam Cerrado oxisols of adequate fertility (corn and cotton plots, prior cotton or soybean residue; soybean plot pH 5.5, base saturation 72%). All trials received full conventional fertilization. A common pre-plant blend of 1500 kg per hectare of an 00-12-15 formulation (with calcium, sulfur, and micronutrients) was applied to every plot. Corn additionally received 400 kg per hectare of urea (about 184 kg N per hectare) split at V2 and V4; cotton received 550 kg per hectare of urea (about 253 kg N per hectare) split across the cycle; soybean received no nitrogen, consistent with reliance on the *Bradyrhizobium* symbiosis, which supplies the bulk of the crop’s nitrogen in Brazilian systems (Hungria and colleagues, 2006). The critical design feature for the present question is that bacterial and biostimulant treatments were superimposed on a complete, non-limiting fertilization program.

### 2.3 Experimental designs and treatments

Each trial was a randomized complete-block design with plots of 3.0 by 6.0 m (18 square meters). Corn comprised seven treatments in four blocks: an untreated control and combinations of *Azospirillum brasilense* (Masterfix Gramíneas, seed), *Methylobacterium symbioticum* (Utrisha N, foliar), and a *Bacillus* phosphorus solubilizer (Omsugo P). Cotton comprised five treatments in four blocks: an untreated control and a biostimulant background (Stimulate plus Hold) with *Methylobacterium symbioticum* (BlueN, foliar) applied at three timings (35, 70, and 90 days after emergence). Soybean comprised four treatments in six blocks: *Bradyrhizobium* spp. (Masterfix L Soja, seed) at two doses, each with and without foliar *Methylobacterium symbioticum* (Utrisha N) at R1; soybean therefore contained no zero-inoculant control, all plots receiving the standard rhizobial inoculation. Product identities, doses, and application timings followed label and program practice and are reported in full in the supplementary material.

### 2.4 Response variables

The primary response was grain or fiber yield: grain yield for corn (sacks per hectare) and soybean (sacks per hectare), and seed cotton and lint yield for cotton (arrobas per hectare). Secondary variables, recorded but treated as supporting, included plant stand, vigor, plant height, biometric yield components, thousand-grain mass, normalized difference vegetation index (GreenSeeker), and, for cotton, High Volume Instrument fiber quality. Leaf nutrient concentrations were determined for soybean at 44 and 65 days after emergence; because these were obtained as a single bulked composite sample per treatment rather than per plot, foliar nutrient data are unreplicated and are reported descriptively only, without inferential testing.

### 2.5 Statistical analysis

Replicate-level data were reanalyzed in Python. For each crop and primary response we fit the randomized complete-block model (treatment plus block) by ordinary least squares and obtained the treatment F test by type II analysis of variance, and separately fit a linear mixed model with treatment as a fixed effect and block as a random effect by restricted maximum likelihood, testing the treatment terms jointly. We report partial eta squared and Cohen’s f (Cohen, 1988) as effect sizes and the coefficient of variation as a measure of experimental precision. Statistical power was summarized as the minimum detectable difference between two treatment means at a significance level of 0.05 and power of 0.80, computed from the residual mean square and replication and expressed as a percentage of the control mean. Equivalence of each active treatment to the untreated control was assessed by two one-sided tests (TOST; Schuirmann, 1987; Lakens, 2017), with equivalence margins set at plus or minus 10% and plus or minus 15% of the control mean and equivalence declared when the 90% confidence interval of the difference fell entirely within the margin. Means reproduced the original trial reports exactly, confirming data fidelity.

## 3. Results

### 3.1 No significant yield response in any crop

No treatment produced a statistically significant yield difference from the control in any crop, and fixed-effect and mixed-model analyses were concordant (Table 2; Figure 1). For corn grain yield, F(6,18) = 0.48, p = 0.82 (mixed model p = 0.82); for cotton seed yield, F(4,12) = 0.03, p = 0.998 (p = 0.998); for cotton lint yield, F(4,12) = 0.13, p = 0.97 (p = 0.97); and for soybean grain yield, F(3,14) = 0.08, p = 0.97 (p = 0.94). Effect sizes were small in cotton (Cohen’s f 0.10 to 0.21) and soybean (0.13) and moderate but non-significant in corn (0.40), where treatment means rose numerically from 111.8 sacks per hectare in the control to 121.8 in the full three-way combination, a non-significant 8.9% increment. Coefficients of variation were 11.9% (corn), 12.3% (soybean), and 6.2 to 6.5% (cotton), the last indicating a notably precise experiment.

**Table 1.**
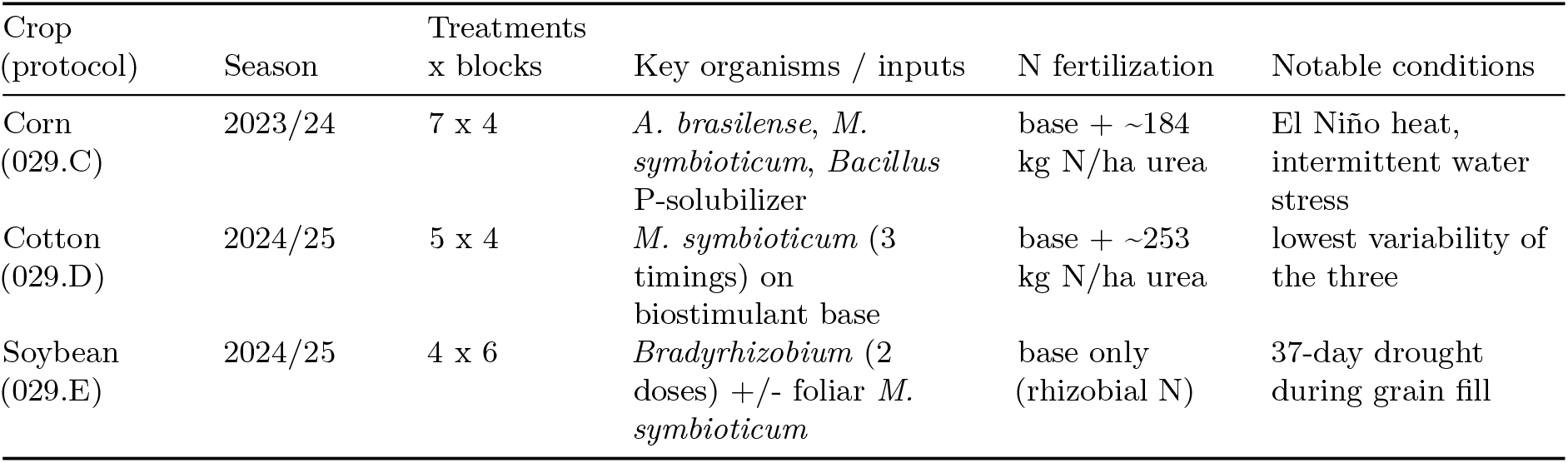
Summary of the three randomized complete-block trials.

| Crop (protocol) | Season | Treatments x blocks | Key organisms / inputs | N fertilization | Notable conditions |
| --- | --- | --- | --- | --- | --- |
| Corn (029.C) | 2023/24 | 7 x 4 | *A. brasilense*, *M. symbioticum*, *Bacillus* P-solubilizer | base + ~184 kg N/ha urea | El Niño heat, intermittent water stress |
| Cotton (029.D) | 2024/25 | 5 x 4 | *M. symbioticum* (3 timings) on biostimulant base | base + ~253 kg N/ha urea | lowest variability of the three |
| Soybean (029.E) | 2024/25 | 4 x 6 | *Bradyrhizobium* (2 doses) +/- foliar *M. symbioticum* | base only (rhizobial N) | 37-day drought during grain fill |

**Table 2.**
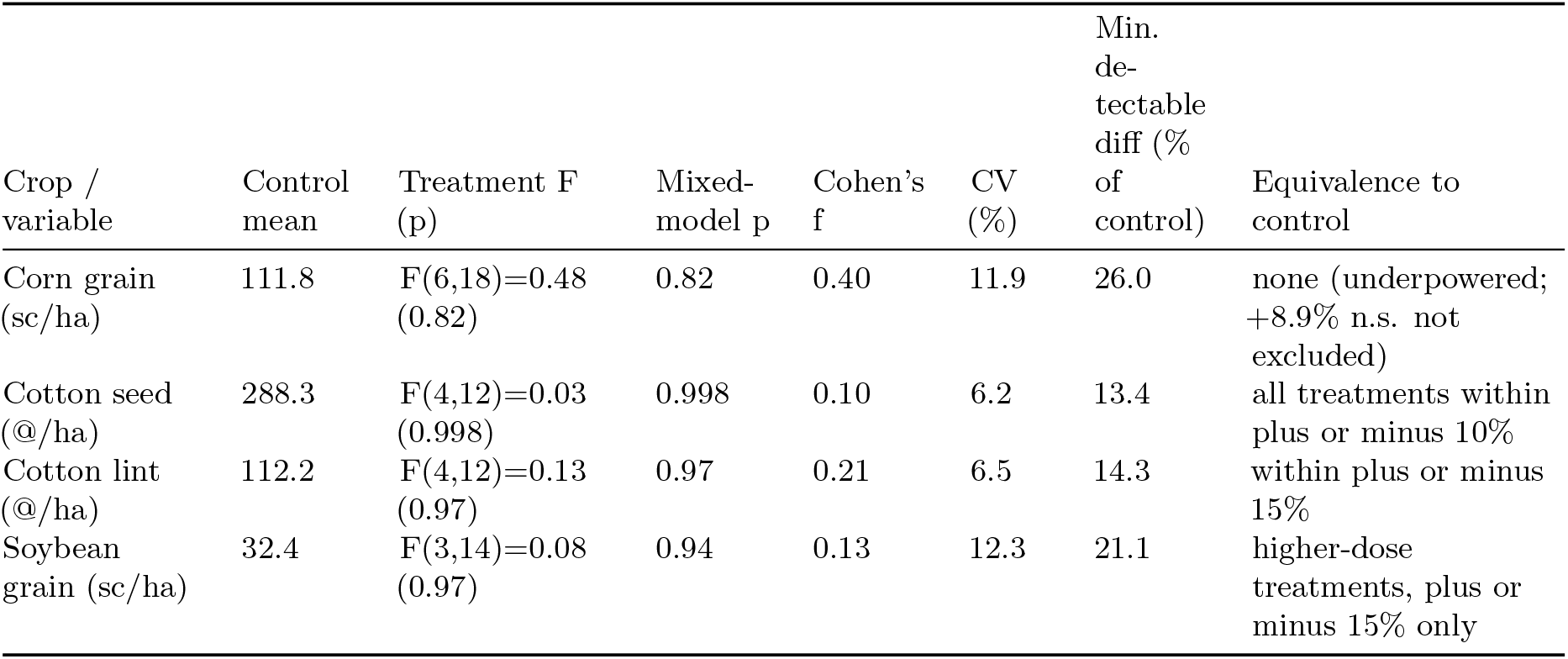
Primary yield results and inference by crop.

**Figure 1.**
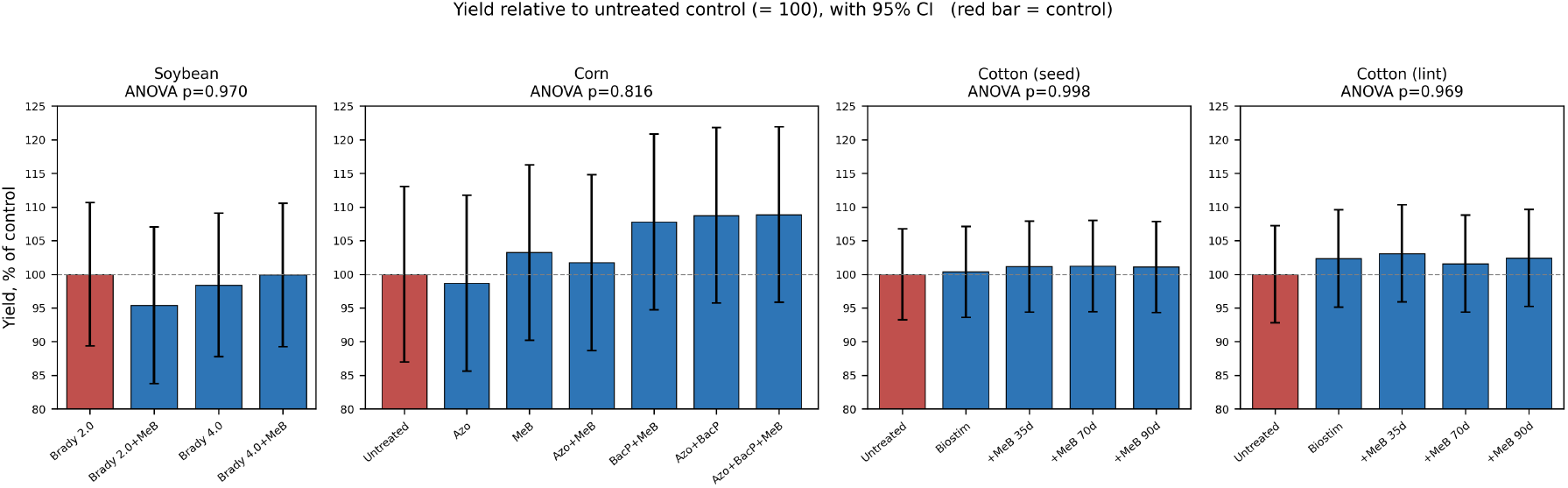
Yield of each treatment relative to the untreated control (set to 100) for the four primary response variables, with 95% confidence intervals. Control bars are shown in red. Treatment effects were non-significant in every crop (analysis of variance p values shown).

### 3.2 Power and the limits of non-significance

The minimum detectable difference at 80% power was 13.4% of the control mean for cotton seed yield and 14.3% for cotton lint, but 21.1% for soybean and 26.0% for corn (Figure 3). In other words, the corn and soybean trials could not have detected differences smaller than roughly one fifth to one quarter of yield, far larger than the single-digit gains (typically 3 to 8%) claimed for these products. Non-significance in corn and soybean therefore cannot be interpreted as evidence of no effect; the experiments lacked the resolution to test the claim.

**Figure 2.**
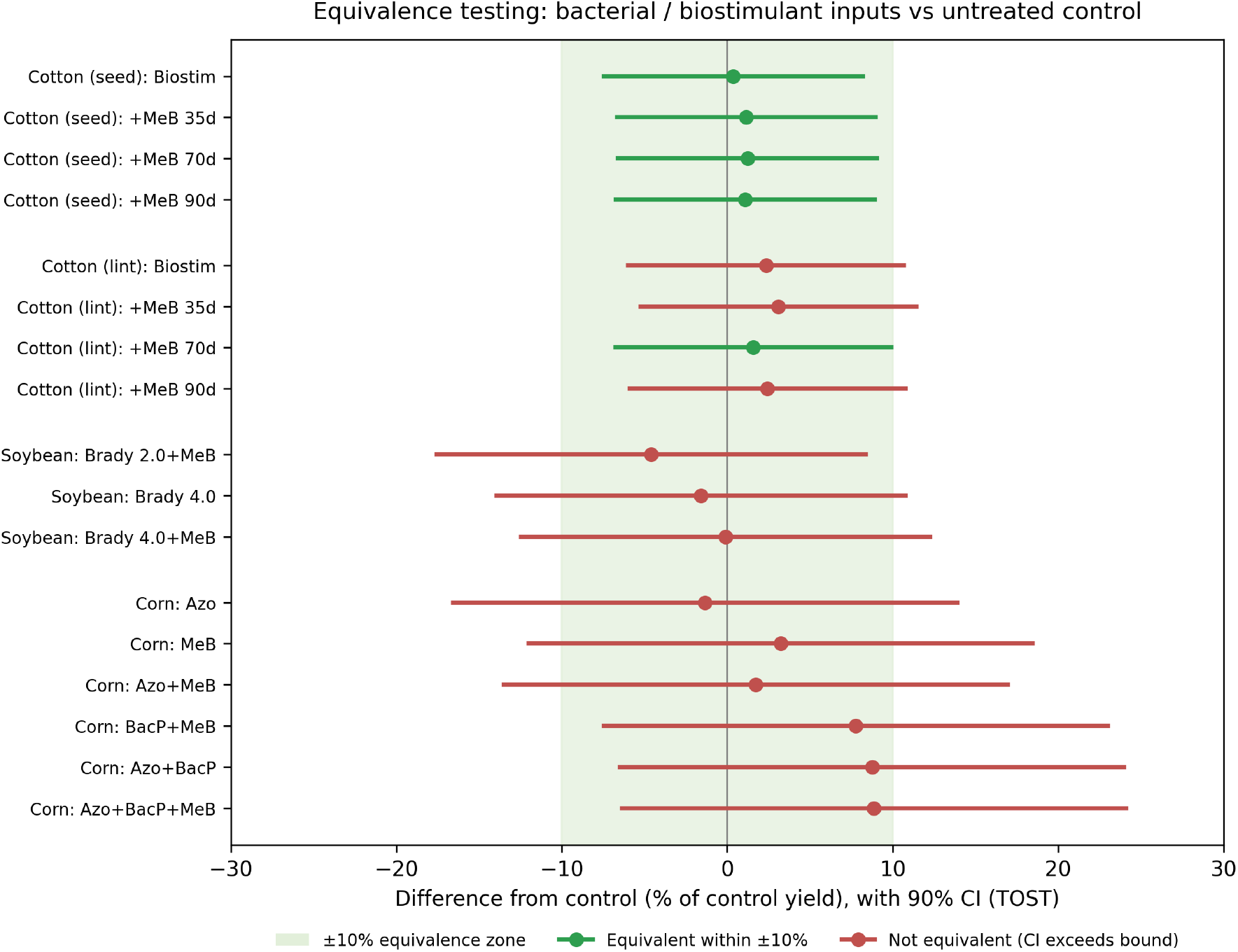
Equivalence testing of bacterial and biostimulant inputs against the untreated control. Points are the difference from control (percentage of control yield) with 90% confidence intervals (two one-sided tests). The shaded band is the plus or minus 10% equivalence zone; green denotes treatments statistically equivalent to control within that zone, red denotes intervals that exceed it. Cotton treatments are equivalent within 10%; corn intervals are wide and inconclusive.

**Figure 3.**
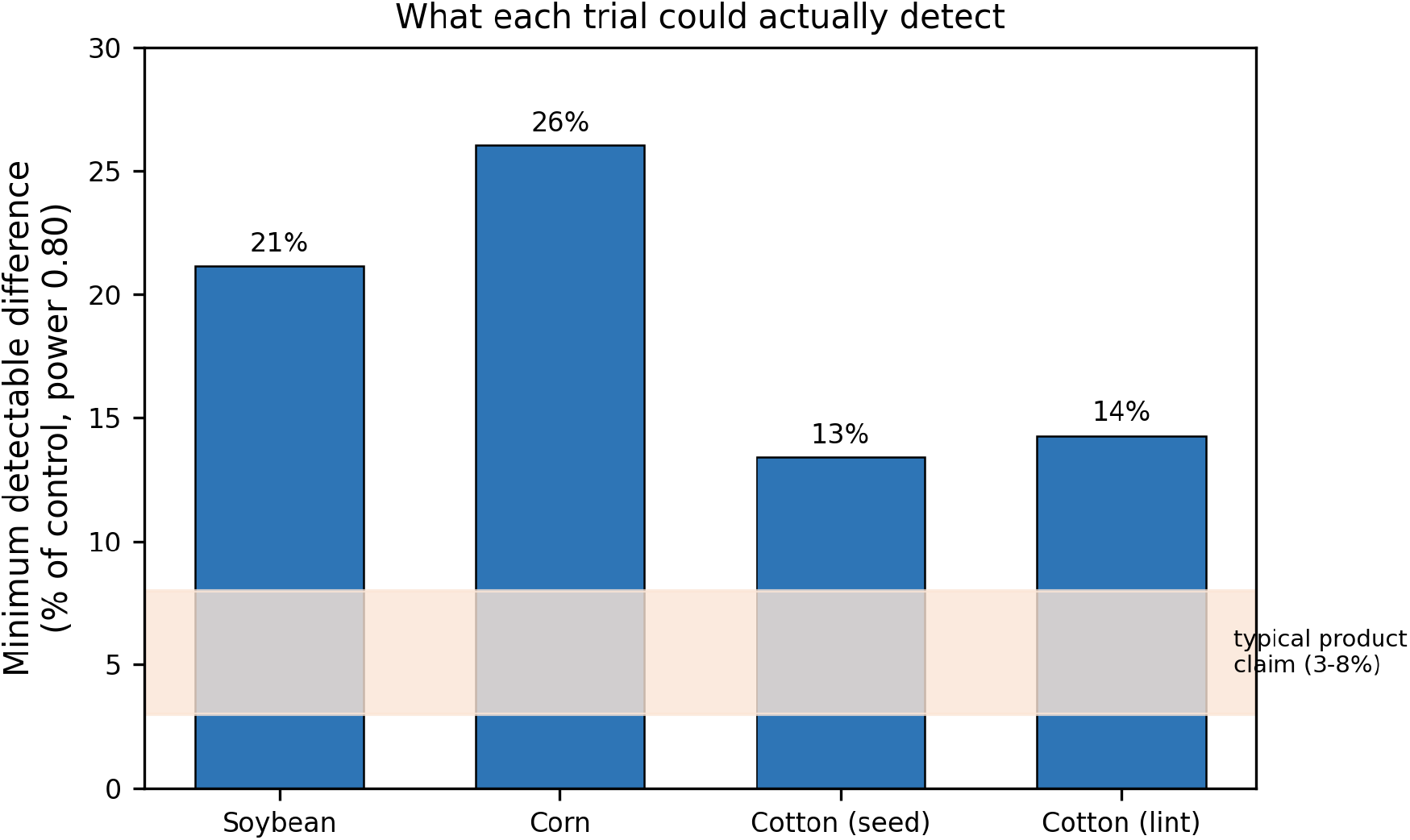
Minimum detectable difference at 80% power for each trial, expressed as a percentage of the control mean. The shaded band marks the 3 to 8% range of yield gains typically claimed for these products; only the cotton trials approach the resolution needed to test such claims.

### 3.3 Equivalence testing

Equivalence testing resolved the asymmetry (Figure 2). In cotton, every treatment was statistically equivalent to the untreated control within a 10% margin for seed yield (TOST p less than 0.035 for all treatments) and within a 15% margin for lint yield; this is an affirmative negative result, a positive demonstration that the inputs added no agronomically meaningful value in the best-powered trial. In soybean, only the two higher-dose *Bradyrhizobium* treatments reached equivalence, and only at the wider 15% margin. In corn, no treatment met equivalence at either margin: the confidence intervals were too wide, and the non-significant numerical gains of 7.8 to 8.9% in the phosphorus-solubilizer and methylotroph combinations could not be statistically excluded. Corn is thus best described as inconclusive rather than null.

### 3.4 Foliar nutrient status (descriptive)

Soybean leaf nutrient concentrations at 44 and 65 days after emergence fell within published sufficiency ranges (Novais and Staut, 2013) for all treatments and showed no enrichment attributable to inoculation. At 65 days after emergence, leaf nitrogen was numerically lowest in the treatment combining *Bradyrhizobium* with the foliar methylotroph (45.9 g per kg, near the lower sufficiency bound) and higher in treatments without the methylotroph (48.0 to 48.2 g per kg). Because these values derive from a single composite sample per treatment, they are not statistically testable and are reported only as an observation consistent with the absence of a measurable nitrogen contribution from the foliar methylotroph; they should be regarded as hypothesis generating.

## 4. Discussion

Across three crops, multiple bacterial chassis, and both seed and foliar delivery, commercial nitrogen-fixing and biostimulant inputs produced no statistically significant yield gain over the control under full conventional fertilization. The contribution of this reanalysis is to move beyond that familiar non-result and quantify what it does and does not support. Where the experiment was adequately powered, in cotton, the answer is unambiguous and affirmative: the inputs were statistically equivalent to the untreated control within 10%, meaning the data positively exclude an agronomically meaningful benefit, not merely fail to detect one. Where the experiments were underpowered, in corn and soybean, intellectual honesty requires the opposite restraint: a minimum detectable difference of 21 to 26% means these trials were never in a position to detect the single-digit gains the products claim, and the non-significant numerical increase of nearly 9% in the corn phosphorus-solubilizer and methylotroph combinations cannot be dismissed. Distinguishing affirmative equivalence from mere non-significance is the central methodological point, and it is one that on-farm evaluations rarely make.

These results sit comfortably within the broader literature once that literature is read carefully. Benefits of *Azospirillum brasilense* in maize are real but are realized predominantly through root stimulation and under partial nitrogen substitution, supporting fertilizer reductions of 20 to 30% rather than additive gains on top of full fertilization (Barbosa and colleagues, 2022). Independent evaluations of *Methylobacterium symbioticum* have likewise found limited and inconsistent field effects, with estimated fixation small relative to crop nitrogen demand (Rodrigues and colleagues, 2024). Our trials test the specific case of inputs superimposed on a complete fertilization program, the very condition under which the literature would predict the smallest marginal value, and that is what we observe. The descriptive foliar nitrogen data, while not inferential, are at least consistent with this reading: the methylotroph treatments showed no leaf nitrogen enrichment.

The framing of this study as a question about added value over standard fertilization is deliberate, and it bounds the conclusions. These trials do not test whether the products can substitute for part of the nitrogen budget, the scenario in which associative inoculants have shown value, because no reduced-nitrogen treatments were included. The relevant and economically consequential question of nitrogen replacement remains open and is the appropriate target of future, suitably designed and adequately powered trials.

Placed against the engineered-fixation frontier, the results carry a clear implication. Programs to install plant-controlled, regulated nitrogenase activity in cereal-associated bacteria aim explicitly to deliver agronomically meaningful nitrogen fluxes that present products do not reliably provide (Ryu and colleagues, 2020; Haskett and colleagues, 2022). Our data quantify how low the current empirical bar sits in commercial fields under standard management: a product equivalent to an untreated control within 10%, as in cotton, contributes nothing an engineered system would need to merely match. The agronomic threshold for a genuinely useful biological nitrogen technology is therefore both well defined and, on this evidence, not met by current associative and methylotrophic inoculants applied over adequate fertilization. This is, we suggest, the more productive way to read a null result of this kind: not as a verdict on a single product line, but as a calibration of the gap that better, more controllable biology will need to close.

Several limitations bound the interpretation and are stated plainly. The full-fertilization background, while it is the framed research question, prevents any inference about nitrogen replacement. Each crop was evaluated at a single site in a single season, and two of the three seasons carried appreciable abiotic stress that depressed yield and widened confidence intervals, contributing to the low power in corn and soybean. The three trials differed in treatment structure and did not share a common control, so they are best read as three coordinated tests of a single question rather than as a factorial whole; soybean in particular lacked a zero-inoculant control. Finally, the foliar nutrient data were unreplicated and can support only description. None of these limitations affects the central, adequately powered finding in cotton, and all of them are addressable in a purpose-designed successor trial that includes reduced-nitrogen treatments, replicated tissue sampling, and multi-environment replication.

## 5. Conclusions

Under full conventional fertilization, commercial nitrogen-fixing and biostimulant inputs added no detectable agronomic value across corn, cotton, and soybean grown at commercial scale in the Brazilian Cerrado. In the best-powered trial the inputs were statistically equivalent to an untreated control within 10%, an affirmative negative result; in the two lower-powered trials, non-significance reflects insufficient resolution rather than demonstrated absence of effect, and a near-9% numerical response in corn warrants better-powered testing. The pairing of equivalence testing with explicit power analysis allows agronomic field data to make calibrated claims that conventional significance testing cannot. Beyond the specific products, the results define the modest empirical performance that current associative and methylotrophic inoculants achieve in the field under standard management, and thereby the threshold that engineered, plant-controlled nitrogen fixation will need to exceed to deliver real agronomic value.

## Data availability

Replicate-level data and the complete analysis code are openly available on Zenodo (https://doi.org/10.5281/zenodo.20696318) and as supplementary material.

## Author contributions

V.H. conceived the reanalysis, co-designed and conducted the field trials, performed the statistical analysis, produced the figures, and wrote the manuscript. L.Y. co-designed and conducted the field trials, curated the primary data, and reviewed the manuscript. Both authors approved the final version.

## Acknowledgments

The authors thank the field and laboratory staff of Cultivo Pesquisa e Desenvolvimento at Fazenda Acalanto.

