## Supplemental Tables S1 S2 S3 S4 S5 for "Do commercial nitrogen-fixing and biostimulant inputs add agronomic value over standard fertilization? An equivalence-based re-analysis of three randomized field trials in the Brazilian Cerrado"

Replicate-level data and analysis code for all tables below are available on Zenodo (https://doi.org/10.5281/zenodo.20696318).

---

#### Table S1. Treatments, products, active organisms, doses, and application timing

##### Corn (029.C, 2023/24), 7 treatments x 4 blocks. Control = untreated.

| Trt | Composition | Active organism(s) | Dose | Timing |
|---|---|---|---|---|
| 1 | Untreated control | none | - | - |
| 2 | Masterfix Gramíneas | *Azospirillum brasilense* | 0.2 L/ha | seed (A) |
| 3 | Utrisha N | *Methylobacterium symbioticum* | 0.333 kg/ha | foliar (22, 40 DAE) |
| 4 | Masterfix + Utrisha N | *A. brasilense* + *M. symbioticum* | 0.2 + 0.333 | seed + foliar |
| 5 | Omsugo P + Utrisha N | *Bacillus* P-solubilizer + *M. symbioticum* | 0.15 + 0.333 | seed + foliar |
| 6 | Masterfix + Omsugo P | *A. brasilense* + *Bacillus* | 0.2 + 0.15 | seed |
| 7 | Masterfix + Omsugo P + Utrisha N | all three | 0.2 + 0.15 + 0.333 | seed + foliar |

**Cotton (029.D, 2024/25), 5 treatments x 4 blocks. Control = untreated.** All treated plots received a Stimulate + Hold biostimulant background.

| Trt | Composition | Active organism | BlueN timing |
|---|---|---|---|
| 1 | Untreated control | none | - |
| 2 | Stimulate + Hold | biostimulant only | - |
| 3 | + BlueN | *M. symbioticum* | 35 DAE |
| 4 | + BlueN | *M. symbioticum* | 70 DAE |
| 5 | + BlueN | *M. symbioticum* | 90 DAE |

##### Soybean (029.E, 2024/25), 4 treatments x 6 blocks. No zero-inoculant control.

| Trt | Composition | Active organism(s) | Dose | Timing |
|---|---|---|---|---|
| 1 | Masterfix L Soja 2.0 | *Bradyrhizobium* spp. | 2.0 mL/kg seed | seed |
| 2 | Masterfix 2.0 + Utrisha N | *Bradyrhizobium* + *M. symbioticum* | 2.0 + 0.33 L/ha | seed + foliar (R1) |
| 3 | Masterfix L Soja 4.0 | *Bradyrhizobium* spp. | 4.0 mL/kg seed | seed |

| Trt | Composition | Active organism(s) | Dose | Timing |
|---|---|---|---|---|
| 4 | Masterfix 4.0 + Utrisha N | *Bradyrhizobium* + *M. symbioticum* | 4.0 + 0.33 L/ha | seed + foliar (R1) |

Product identities: Masterfix Gramíneas and Masterfix L Soja (Stoller); Utrisha N and BlueN (Corteva, both *Methylobacterium symbioticum*); Omsugo P (*Bacillus* phosphorus solubilizer).

---

### Table S2. Complete equivalence (TOST) results for primary yield, every treatment vs control

90% confidence interval of the difference and two one-sided test p values at plus or minus 10% and plus or minus 15% of the control mean.

| Crop | Treatment | Diff (%) | 90% CI (%) | Equiv +/-10% (TOST p) | Equiv +/-15% (TOST p) |
|---|---|---|---|---|---|
| Cotton seed | Biostim | +0.4 | [-7.5, +8.2] | yes (0.024) | yes (0.003) |
| Cotton seed | +MeB 35d | +1.1 | [-6.7, +9.0] | yes (0.033) | yes (0.004) |
| Cotton seed | +MeB 70d | +1.2 | [-6.6, +9.0] | yes (0.034) | yes (0.004) |
| Cotton seed | +MeB 90d | +1.1 | [-6.7, +8.9] | yes (0.032) | yes (0.004) |
| Cotton lint | Biostim | +2.3 | [-6.0, +10.7] | no (0.064) | yes (0.010) |
| Cotton lint | +MeB 35d | +3.1 | [-5.3, +11.4] | no (0.083) | yes (0.013) |
| Cotton lint | +MeB 70d | +1.6 | [-6.8, +9.9] | yes (0.049) | yes (0.007) |
| Cotton lint | +MeB 90d | +2.4 | [-5.9, +10.8] | no (0.066) | yes (0.010) |
| Soybean grain | Brady 2.0 + MeB | -4.6 | [-17.6, +8.4] | no (0.238) | no (0.090) |
| Soybean grain | Brady 4.0 | -1.6 | [-14.0, +10.8] | no (0.125) | yes (0.038) |
| Soybean grain | Brady 4.0 + MeB | -0.1 | [-12.5, +12.3] | no (0.090) | yes (0.026) |
| Corn grain | Azo | -1.3 | [-16.6, +13.9] | no (0.169) | no (0.069) |
| Corn grain | MeB | +3.2 | [-12.0, +18.5] | no (0.225) | no (0.098) |
| Corn grain | Azo + MeB | +1.7 | [-13.5, +17.0] | no (0.180) | no (0.074) |
| Corn grain | BacP + MeB | +7.8 | [-7.5, +23.0] | no (0.401) | no (0.211) |
| Corn grain | Azo + BacP | +8.8 | [-6.5, +24.0] | no (0.445) | no (0.243) |
| Corn grain | Azo + BacP + MeB | +8.9 | [-6.4, +24.1] | no (0.450) | no (0.247) |

Equivalence is declared when the 90% CI lies entirely within the margin. Cotton seed is equivalent at 10% for all treatments; corn meets no margin and is underpowered.

---

### Table S3. Soybean leaf nutrient concentrations (descriptive; single composite per treatment)

Sampled at the fifth node from the apex; one bulked composite per treatment, therefore not replicated and not statistically tested. EMBRAPA SOJA sufficiency ranges shown for reference.

**Nitrogen (g/kg), the variable of interest:**

| | T1 (Brady 2.0) | T2 (+MeB) | T3 (Brady 4.0) | T4 (Brady 4.0+MeB) | Sufficiency |
|---|---|---|---|---|---|
| 44 DAE | 50.4 | 50.3 | 49.2 | 52.0 | 45.0 to 55.0 |
| 65 DAE | 48.0 | 45.9 | 48.1 | 48.2 | 45.0 to 55.0 |

All treatments remained within the sufficiency range at both samplings. At 65 DAE the methylotroph treatment (T2) was numerically lowest. Full multi-element panels (P, K, Ca, Mg, S, B, Cu, Fe, Mn, Zn) at both samplings are in the repository; all elements fell within or above sufficiency for all treatments, with no enrichment attributable to inoculation.

---

### Table S4. Secondary agronomic variables (summary)

No secondary variable differed significantly among treatments in any crop (randomized complete-block ANOVA, Tukey, $p > 0.05$) except isolated, non-systematic pod-class contrasts in soybean. Treatment-level means and full replicate data are in the repository.

| Crop | Variables evaluated | Outcome |
|---|---|---|
| Corn | Phytotoxicity, stand, vigor, thousand-grain mass | No phytotoxicity; uniform stand; vigor differed only for the full combination at one late timing; thousand-grain mass not significant |
| Cotton | Stand, height, node number, boll number and weight, lint percentage, HVI fiber quality (micronaire, strength, length, uniformity, etc.) | All non-significant; uniform crop; fiber quality unaffected by treatment or BlueN timing |
| Soybean | Stand, vigor, shoot fresh mass, GreenSeeker NDVI, height, node and branch number, pod classes, total grains, first-pod insertion, defoliation, thousand-grain mass | All non-significant except isolated pod-class contrasts with no consistent direction; no treatment improved yield components systematically |

---

### Table S5. Site, season, and fertilization detail

| | Corn | Cotton | Soybean |
|---|---|---|---|
| Season | 2023/24 | 2024/25 | 2024/25 |
| Emergence | 9 Jan 2024 | late Dec 2024 | 29 Dec 2024 |
| Pre-plant blend (all plots) | 1500 kg/ha 00-12-15 (+Ca, S, micros) | same | same |
| Nitrogen topdressing | 400 kg/ha urea (~184 kg N/ha), V2+V4 | 550 kg/ha urea (~253 kg N/ha), 3 splits | none (rhizobial N) |
| Plot size | 18 m2 (3 x 6 m) | 18 m2 | 18 m2 |
| Notable conditions | El Niño heat, water stress | lowest CV of the three | 37-day drought during grain fill |

Site: Fazenda Acalanto, São Desidério, BA (12 deg 57 min 52.6 sec S, 45 deg 58 min 58.8 sec W, 833 m), Cerrado biome. Daily meteorological records for each cycle are in the repository.
